# Recruitment of Scc2/4 to double strand breaks depends on yH2A and DNA end resection

**DOI:** 10.1101/2021.07.09.451733

**Authors:** Martin Scherzer, Fosco Giordano, Maria Solé Ferran, Lena Ström

**Affiliations:** Karolinska Institutet, Department of Cell and Molecular Biology, Stockholm, Sweden; Spanish National Cancer Research centre (CNIO), Melcher Fernandéz Almagro 3, 28029 Madrid, Spain

**Keywords:** Scc2/Scc4, Cohesin, DSB, resection

## Abstract

Homologous recombination (HR) enables cells to overcome the threat of DNA double strand breaks (DSB), allowing for repair without the loss of genetic information. Central to the HR repair process is the *de novo* loading of Cohesin around a DSB by its loader complex Scc2/4. Although Cohesin’s accumulation at these sites has been well studied, the prerequisites leading to Scc2/4 recruitment during the repair process are still elusive. Here we investigate which factors are required for recruitment of Scc2 around DSBs in *Saccharomyces cerevisiae*. To address this question, we combined ChIP-qPCR with a GAL-inducible HO-endonuclease system to generate a site specific DSB *in vivo*. We find that Scc2 recruitment relies on yH2A and Tel1, but as opposed to Cohesin, not on Mec1. We further demonstrate that binding of Scc2 depends on and coincides with DNA end resection. Although affected by the impact on resection, this recruitment of Scc2 is not directly facilitated by the RSC, SWR1 or INO80 complexes. Our results shed light on the intricate DSB repair cascade leading to the recruitment of Scc2/4 and the subsequent loading of Cohesin.

## Introduction

A cornerstone in the maintenance of genomic integrity relies on a cells ability to repair DNA damage. This encompasses an orchestrated response necessary to keep its genetic material intact. DNA double strand breaks (DSBs) pose the most hazardous threat to genomic integrity, rendering repair of the same of vital importance[1]. The repair of DSBs is mediated by two major pathways, comprising non-homologous end joining (NHEJ) and homologous recombination (HR). Whereas NHEJ offers rapid repair based on direct end joining, it correlates with increased risk for erroneous repair. HR on the other hand depends on a suitable repair template and is primarily restricted to the S/G2-phases of the cell cycle[2, 3].

The HR repair pathway relies on proteins of the highly conserved Rad52 epistasis group[4]. Among them are Mre11, Rad50 and Xrs2, the constituents of the MRX complex. Credited with the recognition of DSBs, MRX is recruited to broken DNA ends, initiating early stages of DNA end resection and providing a binding platform for the effector kinase Tel1. Recruitment of Tel1 facilitates checkpoint activation, phosphorylation of histone H2A and prevents further progression through the cell cycle[5]. The absence of NHEJ-directing Ku proteins in postreplicative cells then allows the initiation of long-range DNA end resection, carried out by the exonucleases Dna2 and Exo1[6]. The resulting 3’ overhang ssDNA ends are rapidly bound by RPA and provide a binding substrate for Mec1, which in turn reinforces checkpoint activation[7]. The following replacement of RPA with Rad51 enables search for a repair template and the subsequent synthesis of the lost sequence[8]. The presence of a sister chromatid in S/G2 provides cells with a bona fide template to amend DSBs, with minimized loss of genetic information.

Central to the organization of sister chromatids is the ring-shaped protein complex Cohesin. The core Cohesin complex consists of the Smc1/Smc3 heterodimer ATPase, as well the Kleisin subunit Scc1 and the heat repeat protein Scc3. Capable of entrapping DNA within its ring, Cohesin is loaded onto DNA in early S-phase by the separate loader complex composed of the proteins Scc2 and Scc4 (Scc2/4). Upon acetylation of the Smc3 subunit by Eco1, Cohesin is admitted to tether sister chromatids together, a process referred to as establishment of cohesion[9]. Once loaded, ATP hydrolysis allows Cohesin to relocate from the sites of loading and Scc2/4 itself[10]. This was demonstrated by calibrated ChIP-Seq, where only a weak correlation between Cohesin and Scc2/4 binding was found, unless Cohesin’s ATPase function was impaired[11]. In anaphase, after formation of the mitotic spindle, the Scc1 subunit is cleaved by Separase, allowing the segregation of sister chromatids[12]. As opposed to yeast, where Cohesin keeps the chromosomes cohesive along their entire lengths until anaphase, higher eukaryotes remove the majority of chromosome arm bound Cohesin by Separase-independent means already in prophase, leaving only centrometric Cohesin subjected to cleavage of Scc1[13].

The Cohesin complex was first recognized for its significance in DNA DSB repair[14]. It was later shown, that cells were unable to repair DNA damage if they failed to establish cohesion during S-phase[15]. Subsequent experiments demonstrated, that postreplicative *de novo* loading of Cohesin occurs around the break, spanning a region of about 50kb[16, 17]. In line with its loading in early S-phase for sister chromatid cohesion, this process relies on Scc2/4. It was also shown that NIPBL and Mau2, the human homologues of Scc2 and Scc4 respectively, localize to laser microirradiation induced DNA damage stripes as well as Fok1 generated DSBs[18, 19], however DNA binding at breaks has only formally been demonstrated for Scc2 in yeast[20]. Mounting evidence[21–23] attributes the Cohesin loader with a central role in the DNA damage response, yet prerequisites for its recruitment to DSBs are still elusive, despite numerous factors influencing accumulation of Cohesin at DSBs having been identified[16, 17].

Here we address the requirements for Scc2 recruitment to DNA DSBs in yeast using chromatin immunoprecipitation to assess its binding, in selected genetic mutants. We find that the recruitment is largely driven by Tel1 and phosphorylation of histone H2A, whereas Mec1, contrary to its significance for Cohesin loading, is dispensable for this process. We show that Scc2 binding coincides with DNA end resection, where delayed or accelerated end resection affects Scc2 recruitment in a corresponding manner. We further demonstrate that the RSC, SWR1 and INO80 chromatin remodeling complexes are not directly responsible for its recruitment, but do however affect it, presumably through their impact on end resection. We conclude that DNA end resection, together with γH2A, is a driving factor for Scc2/4 recruitment to DSBs, yet by itself insufficient to facilitate Cohesin loading. These findings provide insight into the sequence of events essential for Scc2 recruitment and Cohesin loading during the DSB repair and suggest a potential Cohesin-independent role for Scc2/4 at DSBs.

## Materials and methods

### Yeast strains and growth conditions

All *S. cerevisiae* strains were of W303 origin (ade2-1 trp1-1 can1-100 leu2-3 his3-11,15 ura3-1 RAD5, GAL, psi+). Yeast extract peptone (YEP) supplemented with 40 μg/ml adenine was used as yeast media, unless otherwise stated. For chromatin immunoprecipitation experiments, cells were grown in YEP media supplemented 2% raffinose at 25°C. Arrest in G2/M was induced by addition of benomyl (381586, Sigma) dissolved in DMSO at a final concentration 8 μg/ml for 3 hours, and break induction achieved by the addition of 2% galactose (final) during indicated time periods. Control samples received water. Where applicable, 3-Indoleacetic acid (Auxin - I3750, Sigma) was dissolved in 100% EtOH and added at a final concentration of 1mM. Doxycycline (D9891, Sigma) was dissolved in 50% EtOH and added at a final concentration of 5μg/ml. Control samples received the respective amount of EtOH. To create null mutants, the gene of interest was replaced with an antibiotic resistance marker through lithium acetate based transformation. Some strains were crossed to obtain desired genotypes. For a complete list of strains see table S1.

### FACS analysis of DNA content

G2/M arrest was confirmed by flowcytometric analysis. In brief, 1ml of cultures were fixed overnight in 70% EtOH. Samples were resuspended in 50mM Tris-HCl pH 7.8 and treated with 1mg/mL of RNAse A (12091021, ThermoFisher) at 37°C overnight. Cells were resuspended in FACS buffer (200mM Tris pH7.5, 211mM NaCl, 78mM MgCl_2_) containing 5μg/ml propidium iodide (P4170, Sigma) and sonicated using a Bioruptor Standard (UCD-200, Diagenode). Samples were analysed on a BD FACSCalibur (BD Biosciences) using the CellQuest Pro software.

### Protein extraction and western blotting

To verify Auxin-mediated degradation of target proteins 4 OD units of cells were collected, washed with water and resuspended in glass bead disruption buffer (20mM Tris-HCl pH 8.0, 10mM MgCl_2_, 1mM EDTA, 5% Glycerol, 0,3M ammonium sulfate) supplemented with 1mM DTT, cOmplete protease inhibitor (Roche) and 1mM PMSF. 0,8g of acid washed glass beads (G4649, Sigma) were added and samples were vortexed on a VXR Basic Vibrax (Fisher Scientific) for lysis. Samples were run on Bolt 4-12% Bis-Tris Plus gels (NW04120BOX, ThermoFisher) before transfer to nitrocellulose membranes (GE10600002, Sigma). Anti-FLAG (F1804, Sigma), anti-cdc11 Antibody (y-415, Santa Cruz), anti-AID tag (CAC-APC004AM-T, 2B Scientific) and anti-HA tag antibody (ab137838, abcam) were used in conjunction with appropriate secondary antibodies from the IRDyes series (Licor) and detected on an Odyssey imaging system (Licor).

### Chromatin immunoprecipitation (ChIP) qPCR

ChIP was performed as described [24], with some modifications. 40 OD units of cells were crosslinked in 1% formaldehyde for 30 minutes at room temperature, washed twice in 1x cold TBS, frozen in liquid nitrogen after resuspension in lysis buffer (50mM Hepes-KOH pH 7.5, 140mM NaCl, 1mM EDTA, 0.1% Na-Deoxycholate, 1% Triton X-100, 1x cOmplete protease inhibitor (Roche), 1mM PMSF) and mechanically lysed using a 6870 freezer/mill (SPEX, CertiPrep). WCEs were sonicated using a Bandelin Sonopuls HD 2070.2 mounted with an MS73 probe, for optimally sized DNA fragments (300-700 bp). The protein of interest was purified by over-night incubation with anti-FLAG (Sigma, F1804) or anti-RFA antibody (Agrisera, AS07 214), coupled to dynabeads protein A (Invitrogen). Samples were then washed successively 2x with lysis buffer, 2x with lysis buffer (360mM NaCl), 2x wash buffer (10mM Tris-HCl pH 8, 250mM LiCl, 1mM EDTA, 0.5% Na-Deoxycholate, 0.5% NP-40) and once with TE-buffer. After elution of samples from the beads in elution buffer (50mM Tris-HCl pH 8, 10mM EDTA, 1% SDS) at 65°C for 15 minutes, crosslinking was reversed for both IP and input samples at 65°C overnight. After 1h RNAse (VWR) and 2h Proteinase K (Sigma) treatment, DNA was purified using a QIAquick PCR Purification Kit (QIAGEN). Analysis of DNA was performed by qPCR using Fast SYBR Green Master Mix (Applied Biosystems) on a 7500 FAST Real Time PCR System (Life technologies). Where applicable, data was normalized to an average of N1 and N2 within the same sample. For a list of primers see table S2.

### Measurement of ssDNA at resected DNA ends

10ml of cells (OD = 0.7) were collected and resuspended in 500μl of extraction buffer (100mM NaCl, 50mM Tris-Hcl pH 8.0, 10mM EDTA, 1%SDS) supplemented with 2μl ß-mercaptoethanol (M6250, Sigma) and 2.5μl of Zymolase 100T (20mg/mL). Cells were lysed for 30 minutes at 37°C followed by 5 minutes at 65°C. 250μl KOAc was added followed by incubation on ice for 20 minutes. The lysate was centrifuged and the supernatant was mixed with 0.2ml of 5M NH4OAc and 1 ml isopropanol. The resulting pellet was dissolved in 100μl of TE and 200μl isopropanol, washed with 80% EtOH and resuspended in 50μl of TE. 10μl of each sample were digested with 10U of AciI (R0551S, NEB) and 10U MseI (R0525S, NEB) in a total reaction volume of 30μl using CutSmart Buffer (27204S, NEB) supplemented with 1μl of Ambion RNAseA (AM2271). The digestion was performed overnight at 37°C. Undigested control samples were treated equally with the omission of restriction enzymes. Concentration was measured and adjusted if necessary. 5 serial dilutions of 1:5 were made for each sample and then quantified using Fast SYBR Green Master Mix (Applied Biosystems) on a 7500 FAST Real Time PCR System (Life technologies). For a list of primers see table S2. The difference in average cycles (ΔCt) between digested and undigested samples was measured and the amount of ssDNA calculated according to[25].

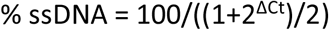

### Pulsed field gel electrophoresis (PFGE)

Break induction at the HO cut site was confirmed with pulsed field gel electrophoresis. The procedure was carried out as previously described[16]. Briefly, cells were collected and fixed overnight in 70% EtOH at −20°C. Samples were resuspended in resuspension buffer (1M Tris base pH7.5, 1.2M Sorbitol, 0.5M EDTA) and lysed in SEMZ buffer (1M Sorbitol, 50mM EDTA, 28mM β-Mercaptoethanol and 2mg/ml Zymolyase 100T (IC320932 VWR)), at 37°C for 90 minutes. Plugs were then prepared with SEZ buffer (1M Sorbitol, 50mM EDTA, 1mg/ml Zymolyase 100T) and 1% low-melting temperature agarose (A9414, Sigma). Embedded cells were then lysed in EST buffer (10mM Tris pH8, 100mM EDTA and 1%Sarcosyl) at 37°C for 45 minutes. After successive equibrilation in 0,5xTBE, plugs were loaded on a 1% PFGE Agarose (1620137, Bio-Rad) gel prepared in 0,5xTBE. Chromosomes were separated on a Biorad Chef DIII (BioRad) at 6 V/cm with a 35,4 - 83,6 second switch time and 120° included angle for 24 hours. Gels were subsequently subjected to southern blot using standard techniques. The PCR product of primers “-1kb Chr VI cut Fw” and “-0,3kb Chr VI cut Rv” was served as probe for the break site. A control probe for chromosome V was generated using primers “Southern Chr V Ctrl Fw” and “Southern Chr V Ctrl Rv” (Table S2).

## Results

### Spatial distribution of Scc2 in the vicinity of a site specific DSB

Cohesin’s dependence on Scc2/4 as a loading factor has been well established, both *in vivo*[26] and *in vitro*[27]. Likewise, it was shown that *de novo* loading of Cohesin at DSBs in postreplicative cells relies on the presence of Scc2 [16, 17]. In agreement with this and fortified by previous Chip-on-chip data in budding yeast, Scc2 can be readily detected at elevated levels around a DSB after 90 minutes of break induction [20], although the requirements for its recruitment are still poorly investigated. We have previously shown that NIPBL, the human homolog of Scc2, localizes to Fok1 generated DSBs through interaction with HP1[18]. However, the absence of an HP1 homolog in *Saccharomyces cerevisiae*[28] raises the question how this recruitment occurs in yeast. To gain insight into this matter, we combined an inducible HO-endonuclease system with ChIP-qPCR to assess the binding of Scc2 around a DSB. An ectopic HO recognition sequence was introduced on Chromosome VI (206,1kb from the left telomere) in equal distance from the centromere and the right telomere, respectively. To make our data comparable between experiments, we utilized previously published low-binding sites of Scc2 on Chromosome V for normalization of the qPCR results[29]. We focused on Scc2, since Scc2 and Scc4 display identical binding patterns *in vivo* in studies analyzing their chromatin association in parallel[10]. The general experimental layout is illustrated in Figure 1A.

**Figure 1.**
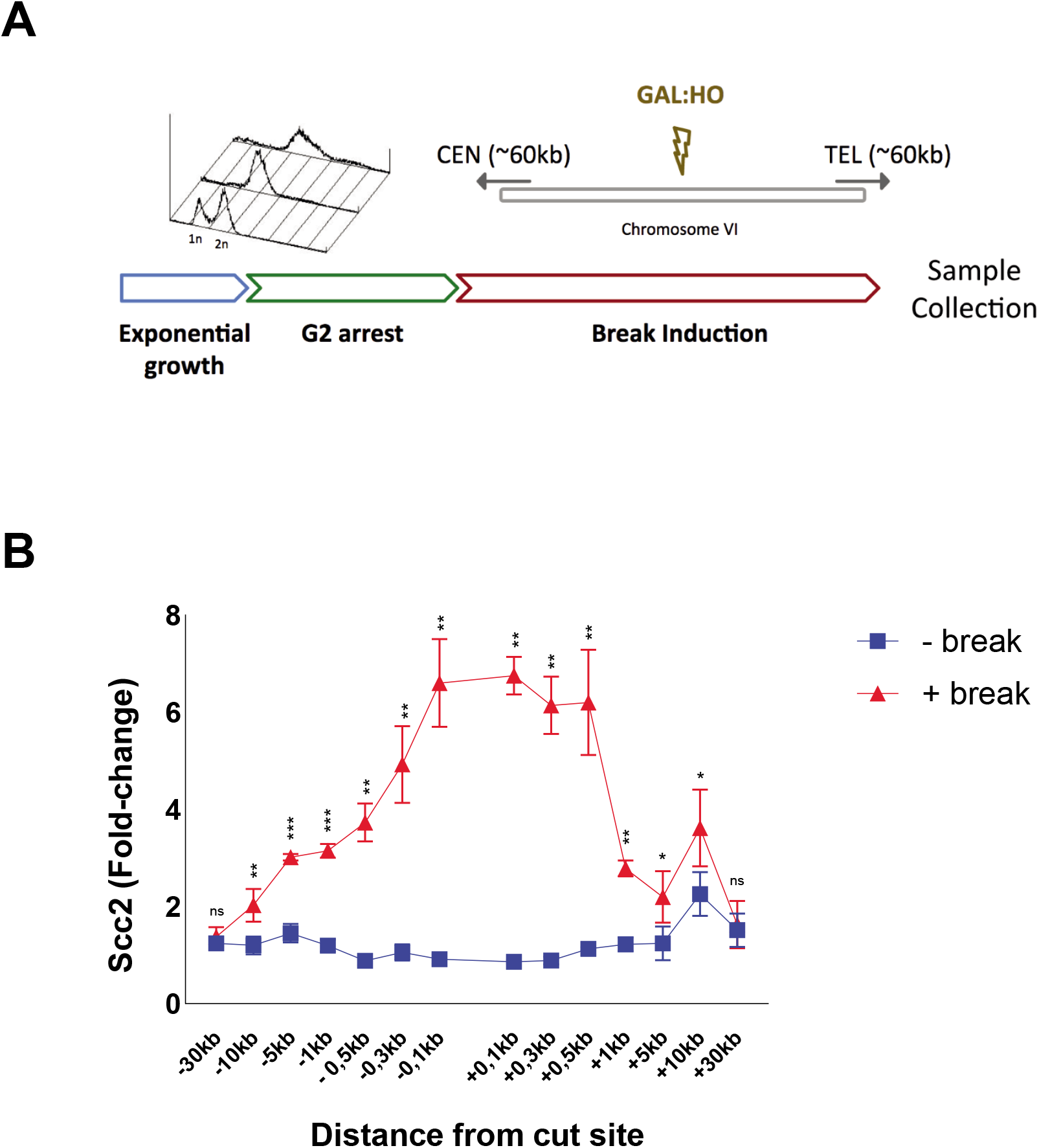
Scc2 is recruited DNA DSBs. **a)** Schematic of the basic experimental setup. Cells in exponential growth phase were arrested in G2 by addition of benomyl for 3h. Galactose was added to induce a DNA DSB on the arm of chromosome 6 in equal distance to the centromere and telomere. Where applicable, cells without induction received water for the same amount of time. After break induction samples were collected and binding of the protein of interest was assessed by ChIP qPCR. Primers used are indicated according to their distance, with “-” referring to upstream and “+” referring to downstream, to the break site. Cells were grown at 25°C throughout. **b)** Scc2 binding is significantly elevated up to 10kb around a DSB, 90 minutes after break induction (red), compared to no break (blue). Data was normalized to low-binding sites. The graph shows means and SD of n =3. Student’s t-test was used to compare +break and -break at respective locations. Significance: *p < 0,05; **p < 0,01; ***p < 0,001; ns = not significant.

Logarithmically growing cells were arrested in G2, followed by addition of either Galactose (+break) to generate a DSB or not (-break) for comparison of Scc2 binding in unchallenged conditions. 90 minutes after break induction samples were collected for ChIP qPCR. At this time point binding of Scc2 increased significantly up to 10 kb around the DSB compared to unchallenged conditions (Fig 1B). This accumulation was comparable on both sides and increasing towards the break site. As opposed to the reported binding pattern of Cohesin at DSBs [16, 17], we found that the most prominent accumulation of Scc2 occurred within 1kb of the DSB. However, since the resolution of ChIP is determined by shearing efficiency (^~^300-700bp), we opted to investigate the requirements for Scc2 binding from 1kb and outwards on the left side of the DSB. The accumulation of Scc2 in a non-mutant strain will henceforth be referred to as wild-type (wt).

### DSB recruitment of Scc2 relies on Tel1 but not Mec1

DSBs are rapidly recognized by the DNA damage sensing Mre11-Rad50-Xrs2 (MRX) complex[30]. Components of the MRX complex have previously been shown to affect the DSB recruitment of yeast and human Cohesin [17, 31]. To investigate the requirement of MRX for Scc2/4 recruitment, we assessed Scc2’s binding in strains lacking either Mre11, Rad50 or Xrs2. Our data demonstrates that binding of Scc2 within 5kb of the DSB was significantly reduced in *mre11Δ* and *rad50Δ* cells, whereas recruitment in *xrs2Δ* was diminished up to 10kb (Fig 2A), rendering the MRX complex an integral part in the recruitment of Scc2. This recruitment did not depend on the nuclease activity of Mre11 as binding was unaffected in deficient backgrounds (Fig S1A).

**Figure 2.**
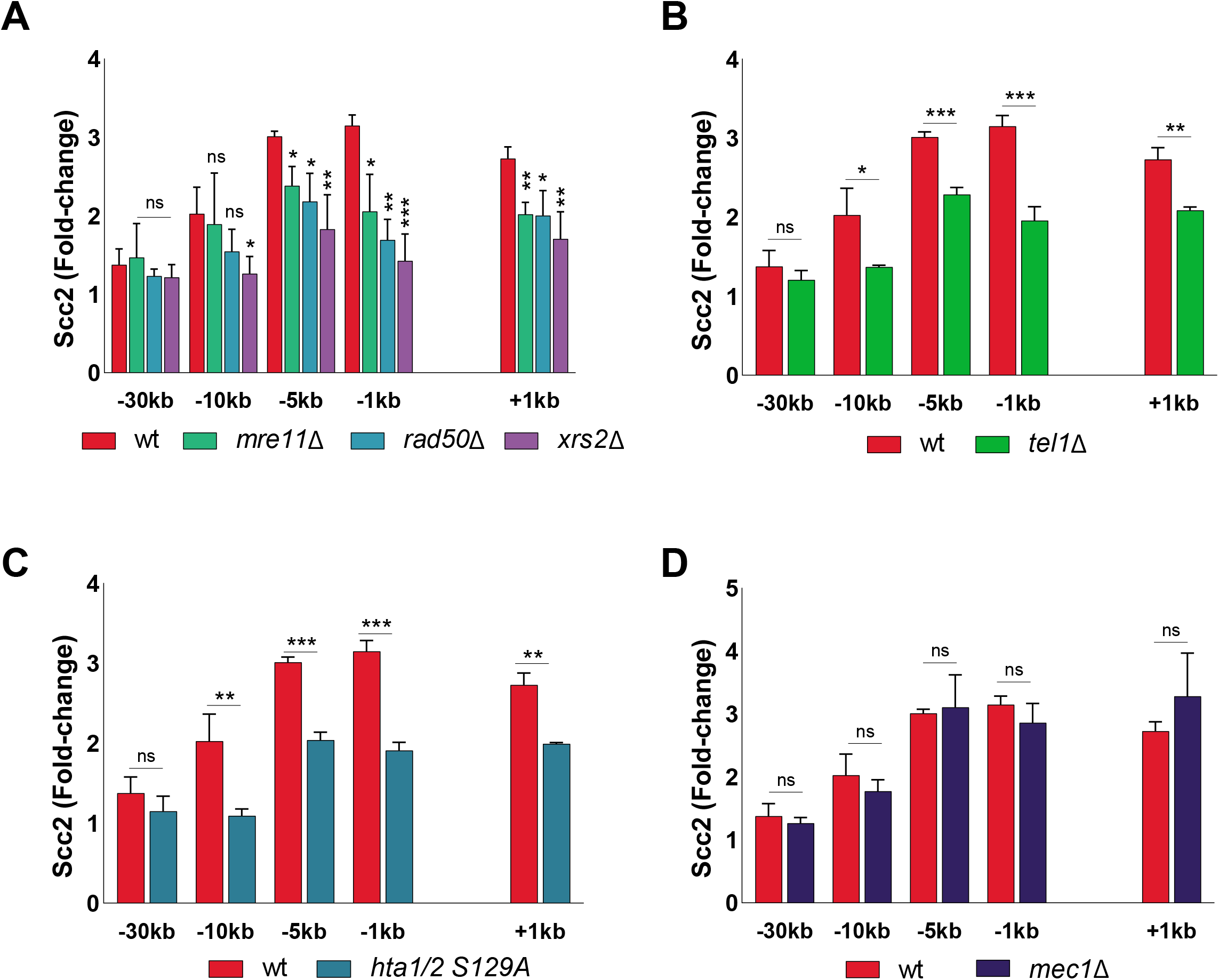
Recruitment of Scc2 depends on the MRX complex, yH2A and Tel1 but not Mec1. ChIP-qPCR of Scc2 binding at the DSB in **(a-d)** wt and strains lacking **a)** Mre11, Rad50 or Xrs, **b)** Tel1 **c)** phosphorylatable alleles of histone H2A or **d)** Mec1. The graphs show means and SD of n =3. Student’s t-test was used to compare normalized values of Scc2 between wt and indicated mutants at respective locations, 90 minutes after break induction. Significance: *p < 0,05; **p < 0,01; ***p < 0,001; ns = not significant.

The initial response to a DSB is accompanied by activation of the DNA damage checkpoint, a process largely regulated by the two kinases Tel1 and Mec1[32]. During checkpoint activation, Tel1 is recruited first, showing a high affinity for broken blunt DNA ends [33], whereas recruitment of Mec1 occurs later[5, 34]. Tel1 was previously reported to be recruited to the DSB by the C-terminus of Xrs2[35]. In agreement with this, deletion of Tel1 led to a considerable reduction in Scc2 recruitment to the break site comparable to the absence of Xrs2 (Fig 2B).

Among the first targets of Tel1 is phosphorylation of histone H2A at Ser129, referred to as γH2A from here on [36]. γH2A has been recognized as an early signal of DNA damage in eukaryotes and is required for the assembly of subsequent effector molecules [37]. In line with previous observations for Cohesin[17], recruitment of Scc2 was indeed impaired (Fig 2C) in strains harboring nonphosphorylateable mutations in both homologs of the H2A gene (hta1-S129A and hta2-S129A)[38].

We next asked, whether absence of Mec1, the second master kinase, augmenting phosphorylation of H2A, would yield similar results as seen for Tel1 (Fig 2B). To our surprise, deletion of Mec1 had no discernable effect on Scc2 recruitment to the DSB (Fig 2D). These results were quite unexpected, as *de novo* loading of Cohesin at the DSB was previously shown to rely on the presence of Mec1, more so than Tel1 [17, 39].

To validate this apparent and potentially interesting discrepancy between Scc2 and Cohesin we wanted to confirm the importance of Mec1 for Cohesin recruitment. In agreement with published results [17, 34], Cohesin failed to be loaded at the DSB in a *mec1Δ* background (Fig 3A). These data highlight different prerequisites for recruitment of Scc2/4 and the subsequent loading of Cohesin.

**Figure 3.**
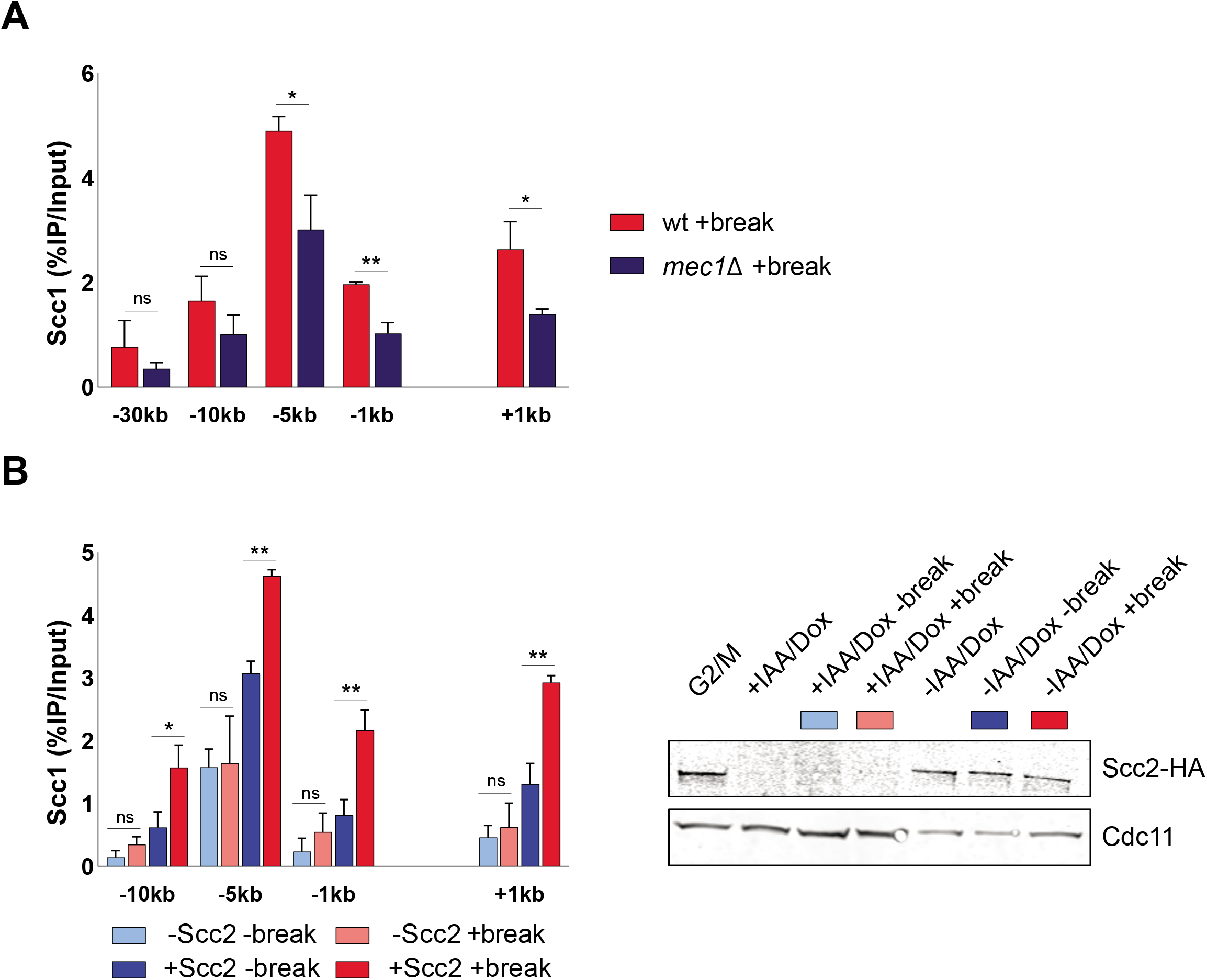
Cohesin relies on Mec1 to be loaded at DSBs by Scc2/4. **a)** ChIP-qPCR of Scc1 binding at the DSB in a wild type and a mec1Δ strain. **b)** Left: ChIP-qPCR of Scc1 binding at a DSB in the presence or absence of Scc2 in an Scc2 degron strain. Cells were grown and arrested as indicated in Figure 1. Prior to break induction, cultures were split in two, with one half receiving Auxin and Doxycycline and the other half receiving a corresponding amount of EtOH for 2 hours to degrade Scc2 or not. Each culture was then split again, totaling 4 and receiving either galactose or water. Right: Western blot showing protein levels of Scc2. Protein samples were taken after 3 hours arrest (G2/M, lane 1), subsequent 2 hours of either IAA/Doxy (+IAA/Dox, lane 2) or EtOH (-IAA/Dox, lane 5) incubation and following 3 hours of either break induction in the presence (-IAA/Dox +break, lane 7) and absence of Scc2 (+IAA/Dox +break, lane 4) or under no break condition in in the presence (-IAA/Dox -break, lane 6) and absence of Scc2 (+IAA/Dox -break, lane 3). Cdc11 served as a loading control. The graphs show means and SD of **a)** n = 3 and **b)** n = 2. Student’s t-test was used to compare values of Scc1 between **a)** wt and mec1Δ or **b)** +break and -break in the presence or absence of Scc2 at respective locations, 180 minutes after break induction. Significance: *p < 0,05; **p < 0,01; ***p < 0,001; ns = not significant.

Since Mec1 recruitment to DSBs relies on RPA-bound single-stranded DNA [7] generated by DNA end resection, this opened for the possibility that Scc2 in itself could be important for the resection process. To test this hypothesis, we generated a strain carrying an auxin-inducible degron (AID[40]) allele of Scc2. Addition of Doxycycline and Auxin after the cells were arrested in G2 led to a substantial reduction in Scc2 protein levels, which remained low during the course of the experiment (Fig 3B). To confirm the lack of Scc2 functionally, we analyzed the accumulation of Cohesin in the presence and absence of a DSB with and without Scc2. Cohesin’s Scc1 subunit could be readily detected on DNA (Fig 3B) and in agreement with our previous results for Scc2 (Fig 1B), this binding was significantly increased in the presence of a DSB up to 10kb around the break. Absence of Scc2 on the other hand resulted in a significant reduction in global Cohesin binding, as has previously been reported to be a consequence of inactivating the Scc2 temperature sensitive allele, *scc2-4*[16, 41]. This was true both for general Cohesin binding in the absence of DNA damage, and *de novo* loading of Cohesin in the presence of a DSB, when compared to control conditions (Fig 3B). Having the possibility to degrade Scc2 efficiently, we next investigated whether Scc2 influenced the degree of resection at the DSB, using a qPCR-based assay adapted from Zierhut et al. [25]. Resection profiles in the presence of Scc2 were comparable to previous studies, reaching around 90% of resected DNA after 6h of break induction 8kb from the DSB[42], whereas resection in a *sgs1Δ/exo1Δ* background was severely impaired (data not shown). Conversely, the absence of Scc2 did not affect the rate of DNA end resection (Fig S2A).

### DNA end resection is a driving factor of Scc2 recruitment

Considering the modest effect of Mec1 on Scc2 recruitment and Scc2 being insignificant for resection, we decided to investigate the importance of resection for the recruitment of Scc2 to DSBs. To monitor the recruitment of Scc2 in the context of DNA end resection we followed the accumulation of Scc2 over a 6h period after break induction, assessing its binding in 90-minute intervals. Ongoing break induction led to a constant increase in Scc2 around the break and elevated levels of Scc2 at −30kb from the break site after 6h (Fig 4A). This increase over time was more prevalent closer to the break. Due to the limitation of the qPCR based approach to measure ssDNA, relying on restriction enzyme cut sites and enzyme efficiency, we instead decided to assess DNA end resection by using RPA ChIP as a readout. Our data shows, that RPA binding (Fig 4B) followed a similar pattern as observed for Scc2, suggesting that recruitment of Scc2 coincides with and might depend on DNA end resection, as has been shown for its binding at stalled replication forks [43]. This prompted us to increase the time of break induction, to allow for DNA end resection. Given the speed of end resection of around 4-5kb/h [44, 45] and the kinetics of break formation, we decided to analyze the recruitment of Scc2 after 180 minutes from here on, as resection proceeds well beyond 10kb (Fig 3B) in the majority of cells at this time. To assess the impact of DNA end resection on Scc2 recruitment, we analyzed its binding in a strain carrying deletions of Sgs1 and Exo1, noted for a severe resection deficiency beyond a few hundred base pairs. Recruitment of Scc2 to the DSB was drastically decreased in the *sgs1Δexo1Δ* background (Fig 4C). Consistent with the impairment of long-range resection, this effect was less prominent proximal to the break (Fig S2B), confirming the significance of end resection for Scc2 recruitment.

**Figure 4.**
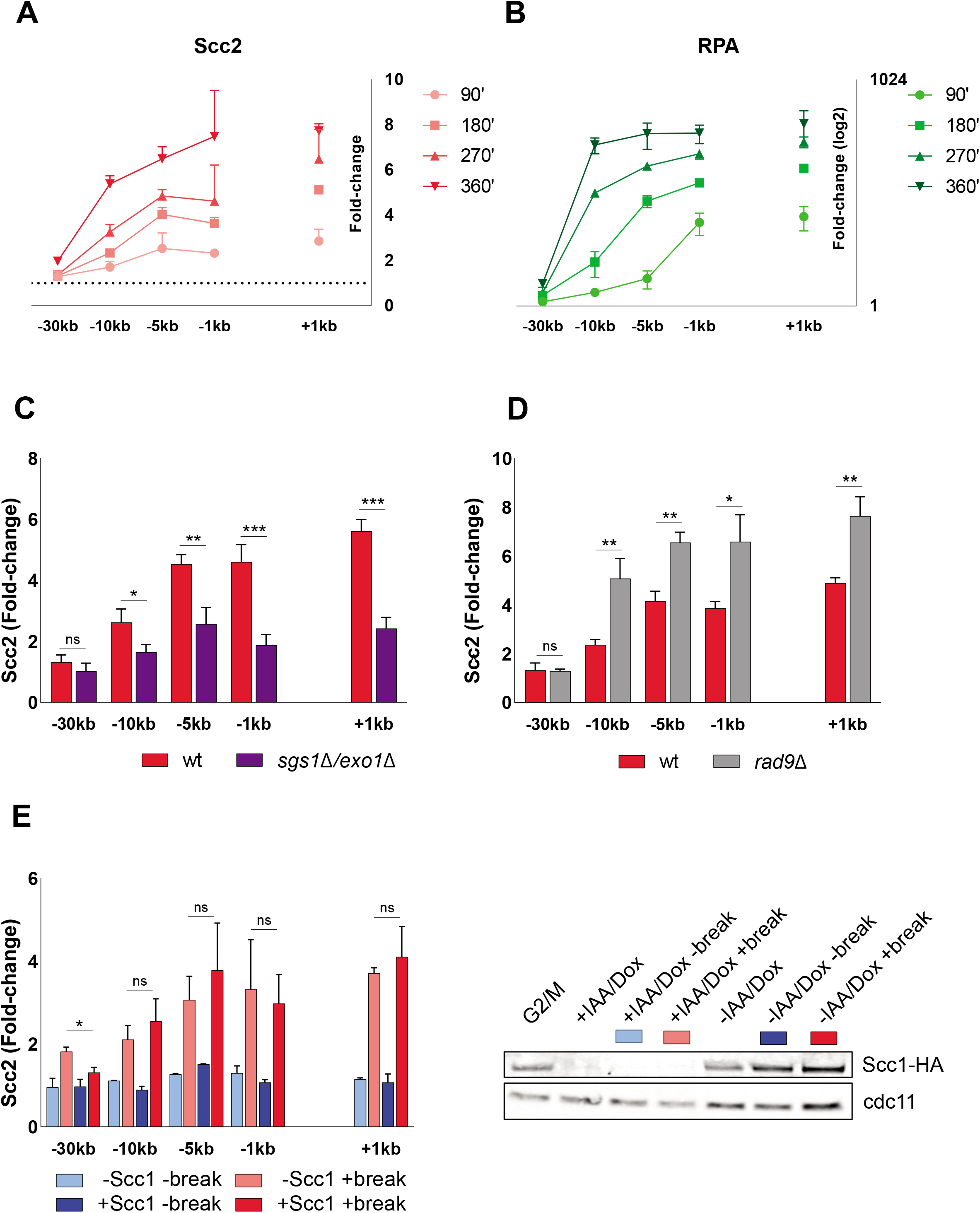
Scc2 binding at DSBs accumulates over time and depends on DNA end resection. ChIP-qPCR time course of **a)** Scc2 and **b)** RPA binding at the DSB. Samples were taken over a 6 hour period at 90 minute intervals. A log2 scale was used to visualize RPA binding. ChIP-qPCR of Scc2 binding at a DSB in **(c&d)** wt and strains lacking **c)** Sgs1 & Exo1 **d)** Rad9. **e)** Left: ChIP-qPCR of Scc2 binding at a DSB in the presence or absence of Scc1 in an Scc1 degron strain. Cells were grown and arrested as indicated in Figure 1. Prior to break induction, cultures were split in two, with one half receiving Auxin and Doxycycline and the other half receiving an equal and appropiate amount of EtOH for 2 hours to degrade Scc1 or not. Each culture was then split again, totaling 4 and receiving either galactose or water. Right: Western blot showing protein levels of Scc1. Protein samples were taken after 3 hours arrest (G2/M, lane 1), subsequent 2 hours of either IAA/Doxy (+IAA/Dox, lane 2) or EtOH (-IAA/Dox, lane 5) incubation and following 3 hours of either break induction in the presence (-IAA/Dox +break, lane 7) and absence of Scc1 (+IAA/Dox +break, lane 4) or under no break condition in in the presence (-IAA/Dox -break, lane 6) and absence of Scc1 (+IAA/Dox -break, lane 3). Cdc11 served as a loading control. Graphs show means and SD of (**a,b & e)** n = 2 and (**c & d)** n = 3. Student’s t-test was used to compare values of Scc2 between wt and **c)** *sgs1Δ /exo1Δ* or **d)** rad9Δ at respective locations, 180 minutes after break induction. In **e)** binding of Scc2 in +break was compared between the presence or absence of Scc1 at respective locations, 180 minutes after break induction. Significance: *p < 0,05; **p < 0,01; ***p < 0,001; ns = not significant.

If resection as such is a determining factor for Scc2 recruitment, then increased resection should augment the Scc2 binding. To test this, we assessed Scc2 recruitment in a strain lacking Rad9 which causes DNA end resection to proceed at an accelerated pace[46]. Consistent with our hypothesis, deletion of Rad9 resulted in significantly elevated levels of Scc2 recruitment to the DSB compared to wild type (Fig 3D). Based on our results, we conclude that DNA end resection is a decisive process for Scc2 recruitment, whereas loading of Cohesin requires additional events to take place, such as recruitment of Mec1 (Fig 3A).

A recent study by Arnould et al[47], investigating the role of topologically associating domains (TADs) in DNA damage repair in human cells, proposed a model where a loop extruding mechanism allows rapid phosphorylation of histone H2AX as DNA is reeled in by DSB anchored Cohesin. Considering these findings, our observation that Scc2 “emanates” from the break site over time could be explained by Cohesin-mediated loop extrusion. To address the possibility that Scc2/4 would load Cohesin at the break site and then be shuttled away by loop extruding Cohesin, causing a technical artifact, we employed a strain carrying an Scc1-AID construct. This allowed for degradation of Scc1, which consequently would interfere with potential Cohesin-mediated loop extrusion at the break. Degradation of Scc1 prior to break induction caused no significant reduction in the recruitment of Scc2 to the DSB (Fig 4E). Interestingly, binding was significantly increased 30kb from the DSB, raising the possibility that yH2A dynamics could also be affected by Cohesin in yeast. A minor, albeit significant, decrease was observed at 5kb in unchallenged conditions. Based on these results we conclude that recruitment of Scc2 on one hand occurs independently of Cohesin and that its “emanation” on the other hand is unaffected by possible Cohesin-mediated loop extrusion at DSBs.

### Impact of chromatin remodeling on Scc2 recruitment

Based on our finding that DNA end resection appears to be a critical factor in the recruitment of Scc2, we next addressed the role of chromatin remodeling. As DNA end resection is tightly regulated by chromatin remodeling[48], we channeled our attention first to the RSC complex. It was previously demonstrated that the Sth1 ATPase subunit of the RSC complex acts as a chromatin receptor, facilitating the binding of Scc2/4 and subsequent loading of Cohesin[49]. Given the significance of the RSC complex for loading of Scc2/4 during an unchallenged cell cycle and its central role in the early processing of DSBs[50], we asked whether Sth1 was equally integral for the recruitment of Scc2 to DSBs. Rather surprisingly, recruitment of Scc2 to DSBs at restrictive temperature remained largely comparable to its recruitment at permissive temperature in a strain harboring the temperature sensitive allele *sth1-3* (Fig 5A). However, recruitment was overall reduced compared to wild-type (compare to Fig 5C, D), also under permissive conditions.

**Figure 5.**
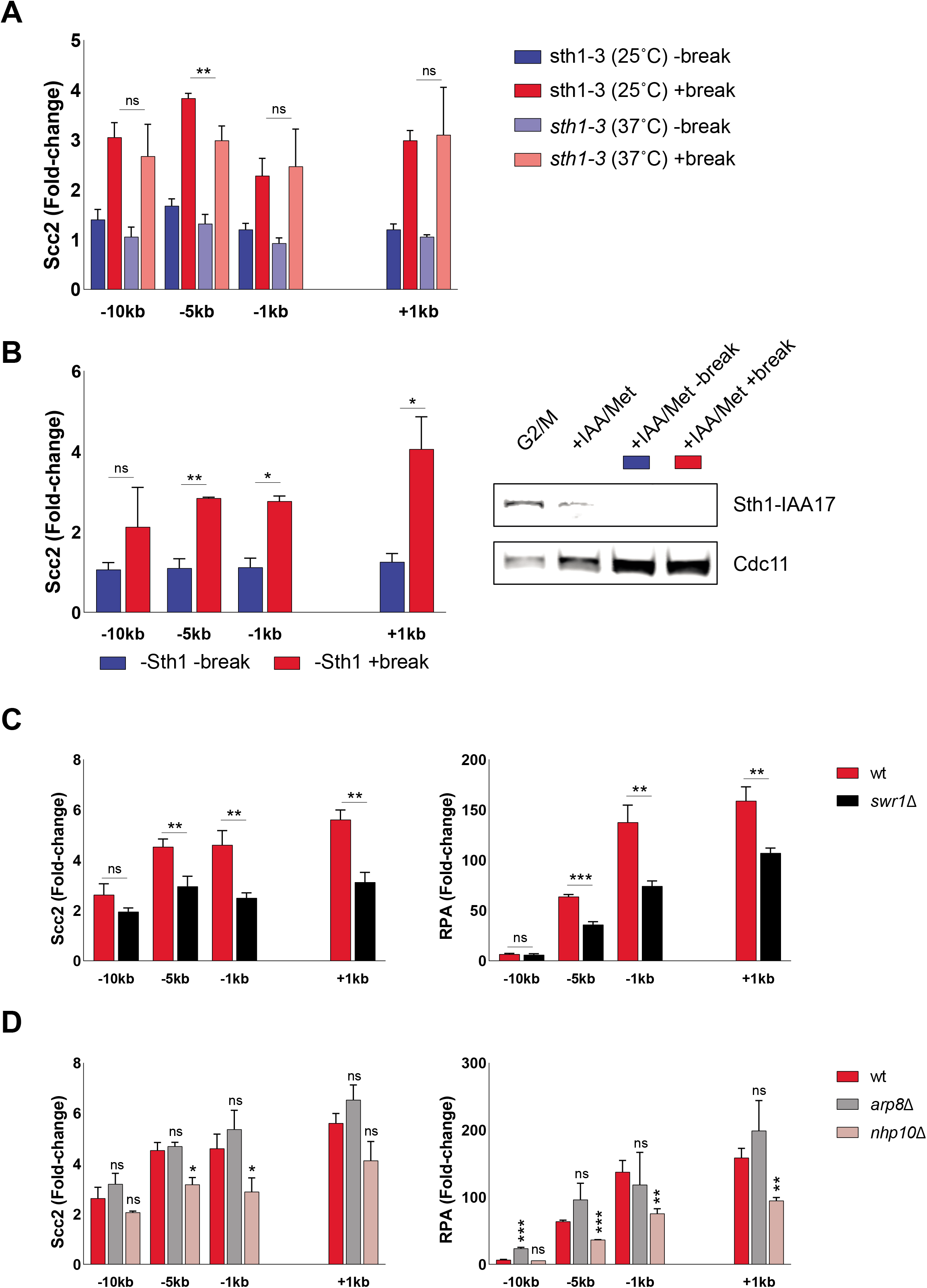
Scc2 recruitment is not directly facilitated by RSC, SWR1 or INO80. **a)** ChIP-qPCR of Scc2 binding at the DSB in an *sth1-3* background. Cells were grown and arrested as indicated in Figure 1. 30 minutes prior to break induction, cultures were split into two with half grown at 25°C and the other half grown at 37°C to inactivate Sth1. Each culture was then split again, totaling 4 and receiving either galactose or water. **b)** Left: ChIP-qPCR of Scc2 binding without Sth1 in the presence of absence of a DSB. Cells were grown in -met media and shifted to benomyl containing YEP media supplemented with 2mM Methionine for 3 hours. Cultures were then split, totaling 2 and receiving either galactose or water. Poor growth prohibited arrest in -met media, denying analysis of suitable control conditions. Right: Western blot showing protein levels of Sth1. Protein samples were taken after 3 hours arrest (G2/M, lane 1), 3 hours after arrest (+IAA/Met, lane 2) and following break induction (+IAA/Met +break, lane 4) or not (+IAA/Met -break, lane 3). Cdc11 served as a loading control. **c)** ChIP-qPCR of Scc2 (left) or RPA (right) binding at a DSB in wt and *swr1Δ*. **d)** ChIP-qPCR of Scc2 (left) or RPA (right) binding at a DSB in wt, *arp8Δ* and *nhp10Δ*. Graphs show means and SD of n =3. Student’s t-test was used to compare normalized values of Scc2 between **a)** +break with and without Sth1 **b)** in the absence of Sth1 with and without break **c & d)** wt and indicated mutants at respective locations, 180 minutes after break induction. **c & d)** Additionally, a student’s t-test was used to compare normalized values of RPA between wt and indicated mutant. Significance: *p < 0,05; **p < 0,01; ***p < 0,001; ns = not significant.

To corroborate these findings, we in addition utilized a strain expressing AID-tagged Sth1 under a repressible MET3 promoter[49]. Presence of methionine and auxin reduced the Sth1 protein level by more than 80%, prior to break induction (Fig 5B). Unfortunately, cells grown under permissive conditions failed to arrest in G2/M due to poor growth in minimal media without methionine, combined with Raffinose as a suboptimal carbon source. Despite the absence of the ideal control, recruitment of Scc2 was significantly increased in the presence compared to the absence of a DSB despite Sth1 being absent (Fig 5B), and comparable to the recruitment in the *sth1-3* strain (Fig 5A). We thereby conclude that the role of the RSC complex in Scc2/4 loading does not extend to the DNA damage repair response. We reason that the reduction in recruitment is rather due to indirect effects on DNA end resection[50].

Next, we asked if other chromatin remodelers could be responsible for the recruitment of Scc2 to DSBs. Considering the significance of yH2A for the recruitment of Scc2 (Fig 2C), the SWR1-C and INO80 chromatin remodeling complexes posed as suitable candidates since both have been shown to be able to bind to yH2A[51, 52]. Deletion of Swr1, the ATPase subunit of SWR1-C, lead to a severe reduction in Scc2 at the DSB (Fig 4C). We reason that this might be due to the resulting hampered incorporation of histone H2AZ, negatively affecting DNA end resection[53]. Although described as resection-proficient, RPA coverage in *swr1Δ* confirmed perturbed generation of ssDNA under these experimental conditions (Fig 5C right graph).

Since the ATPase subunit of INO80 is essential in W303[54], we decided to address its role for the recruitment of Scc2 by using strains harboring deletions of the Arp8 and the Nhp10 subunits, which interfere with the chromatin remodeling ability and the recruitment of INO80 to DSBs respectively[55]. Interestingly, although both mutants are renowned for deficient DNA end resection[56], only deletion of Nhp10 resulted in a significant reduction in the recruitment of Scc2 to the DSB, whereas deletion of Arp8 remained indistinguishable from wild type (Fig 5D, left graph). However, in agreement with the observed binding pattern for Scc2, only *nhp10Δ* resulted in a reduction of RPA filament formation around the break, while *arp8Δ* remained comparable to wild type. This discrepancy has been addressed previously, suggesting that DNA end resection is mildly impeded in *arp8Δ* strains, despite RPA binding remaining unchanged or even slightly increased[57]. Considering RPAs strong affinity for ssDNA[58], we reason that although DNA end resection may be impaired in *arp8Δ*, singlestranded nucleofilaments or regions displaying ssDNA may still be able to form, mimicking DNA end resection.

Overall, our data suggest that recruitment of Scc2/4 to DNA DSBs is mediated through phosphorylation of H2A by Tel1, but as contrary to Cohesin not Mec1, and occurs during DNA end resection. We reason that this recruitment is favored by the generation of ssDNA, which occurs during DNA end resection. Furthermore, Scc2/4 recruitment is not directly facilitated by the chromatin remodeling complexes RSC, INO80 or SWR1 as the effect on recruitment was comparable to the degree of impaired resection.

## Discussion

Cohesin’s accumulation at DNA DSBs and its dependency on Scc2/4 in any context are both well documented. However, research in the field of DNA damage repair has been focusing almost exclusively on Cohesin[59]. In order to get mechanistic insight into how Cohesin is loaded at DSBs it is therefore indispensable to understand how its loader gets there in the first place. Here we provide the first investigation focusing on the recruitment of Scc2 to DNA DSBs in budding yeast. We find that its accumulation depends mainly on γH2A and DNA end resection, neither of which alone suffices to facilitate the recruitment. Although Cohesin and its loader share several factors needed for their accumulation at DSBs, our study also uncovered an unexpected difference between the recruitment of Scc2 and the loading of Cohesin. Whereas both Tel1 and Mec1 are required for *de novo* loading of Cohesin at DSBs, Mec1 is dispensable for the recruitment of Scc2.

The significance of DNA end resection for HR repair is well established, yet only recently its impact on Cohesin is starting to gain traction[43, 60]. We show that Scc2 recruitment emanates from DSBs coincident with ongoing resection. Similar observations have been made for chromatin remodelers modulating DNA end resection[42], however we did not find evidence for Scc2s involvement in this process (Fig S2A). Supported by the fact that Scc2s binding at DSBs was drastically reduced in a *sgs1Δ/exo1Δ* mutant, this points towards an epistatic relationship of Scc2 recruitment and DNA end resection.

In accord with previous data for Cohesin, we find that the MRX complex is required to facilitate the recruitment of Scc2 to DSBs. This dependency most likely relies on MRX’ ability to recruit Tel1 and the resection machinery[61], as recruitment of Scc2 was comparable to wild type in a nuclease-deficient Mre11 D56A & H125A mutants which still allow complex formation[62, 63] (Fig S1A). It was shown that “clean” DSBs, that is breaks without DNA abducts, can bypass the need for the initial incision at DNA ends by MRX and Sae2 to promote Dna2 and Exo1[64], which would also explain, why deletion of Sae2 had no effect on the accumulation of Scc2 at DSBs (Fig S1B).

Most strikingly, we find that deletion of Mec1, has no effect on Scc2 recruitment, yet impairs Cohesin loading at the DSB. The exact nature of Cohesin’s dependency on Mec1 is still unknown. It was shown that phosphorylation of Scc1 at Ser83 by Chk1, presumably downstream of Mec1, was required for the generation of damage induced cohesion, yet loading around the break was unaffected in S83A mutants[39]. Although both Scc2 and Scc4 harbor multiple consensus motifs for Mec1/Tel1[65], we were unable to detect phosphorylation of these sites in response to DNA damage by mass spectrometry (data not shown), dampening the possibility of a direct effect on Scc2/4 by either. *In vitro* experiments have demonstrated that second strand capture of Cohesin is favored if the target is single-stranded. These events were counteracted by addition of RPA[66]. Applying this concept on a DSB, it can be envisioned that Mec1 phosphorylates RPA[67], destabilizing its association with DNA[68] and thereby enabling the loading of Cohesin by Scc2/4. It could also be that recruitment of Mec1 affects the chromatin landscape around the break, as has been observed for stalled replication forks[69], which in turn favors the loading of Cohesin[49].

The requirement of γH2A for the recruitment of Scc2 is consistent with what has been observed for Cohesin[17]. However, previous studies have shown that the *hta1-S129A* background causes accelerated end resection[70], indicating that DNA end resection by itself is insufficient for Scc2 recruitment. Conversely, it was also shown that γH2A spreading increases in the absence of Sgs1/Exo1[71], indicating that also γH2A alone is insufficient for Scc2 recruitment. The fact that recruitment of Scc2 was drastically increased in a *rad9Δ* background, likewise shown to have accelerated resection kinetics but an unaltered γH2A profile[5], lead us to the conclusion that recruitment is not directly mediated by DNA end resection but rather augmented by it.

Due to the complex interplay of DNA end resection and chromatin remodeling, we reasoned that chromatin remodelers could dictate the recruitment of Scc2 depending on the biological context as previous studies have demonstrated to be case for Scc2[49]. Given the role of the RSC complex in the DNA damage response[50] and the requirement of RSC components for Cohesin loading at DSBs[72] we expected similar results for Scc2. Although recruitment was reduced overall compared to genuine wild type cells, we failed to see an effect drastic enough to allow the conclusion that Sth1 serves as an Scc2/4 loading factor at DSBs. We reason that this reduction is rather due to impaired resection kinetics.

Based on our results that Scc2 recruitment depends on γH2A, we then decided to focus on SWR1C and INO80C, both of which have been proposed to depend on γH2A[52, 73], though this claim has been contested[71]. Responsible for the H2AZ metabolism at DSBs, SWR1 is recruited to breaks facilitating the incorporation of H2AZ[74], whereas INO80C catalyzes its removal in addition to general nucleosomal eviction[57]. Although *in vitro* experiments have demonstrated that incorporation of H2AZ benefits DNA end resection, the absence of Swr1 does not cause a resection defect *in vivo*[42, 53]. Seemingly in contrast to our hypothesis, recruitment of Scc2 was markedly reduced in *swr1Δ*. Furthermore, although INO80C mutants have been noted for resection defects, we did not observe a reduction of Scc2 recruitment in *arp8Δ* mutants. To circumvent the substantial amount of conflicting data regarding resection defects in these mutants[57, 73, 75], we decided to measure the generation of ssDNA under our experimental conditions in respective strains. RPA binding agreed with our hypothesis and demonstrated that the RPA filament formation was impaired in *swr1Δ* and *nhp10Δ*, but increased in *arp8Δ*, which could account for the slight, albeit insignificant, increase in Scc2 at the DSB. We hypothesize that the reduction of RPA seen in *swr1Δ* cells could be due to delayed HO kinetics[76], although break induction at the experimental endpoint was comparable to wild type (Fig S3C). In support of this, we failed to induce appreciable DSBs in an *htz1Δ* background within 3 hours (Fig S3D). We want to note here, that INO80Cs mechanism may be altered depending on laboratory strain background. While the INO80 ATPase subunit is essential in W303, it is not so in S288C[54]. Furthermore, deletion of INO80C components have been shown to cause polyploidy in a S288C derivate strain, with Arp8 being a notable exception[77]. However, we observed significant polyploidy in our *arp8Δ* mutant in a W303 background (S3A), which was absent in other strains. In any case, we reached a similar conclusion as with Sth1 and believe that neither SWR1C nor INO80C are directly responsible for recruitment of Scc2 as its binding correlated well with RPA coverage. However, as our investigation did not comprehensively address all chromatin remodelers, we cannot exclude the possibility that other complexes are responsible for Scc2/4 “loading” at DSBs. This would justify a more thorough investigation, beyond the scope of this study.

The exact mechanism that facilitates the recruitment of Scc2 to DSBs remains to be determined. Although ssDNA was shown to be bound poorly by Scc2/4 *in vitro*, its affinity for Y-fork DNA was comparable to dsDNA[27]. It can be envisioned, that in the process of end resection a similar intermediate is formed, favoring its recruitment. We have previously demonstrated that inactivation of Scc2 in yeast modulates transcription globally and in response to a DSB, affecting DSB proximal genes in particular[23]. Studies in human cells have shown that transcriptional repression at DSBs is mediated by NIPBL and Cohesin[78], whereas in yeast this process is mediated by end resection[70]. Accumulating evidence highlights the significance of RNA and transcription in the DNA damage response and the modulation of resection[79]. Considering our data, it would be interesting to address the impact of transcription at DSBs on Scc2 recruitment and vice versa[80].

In summary, we demonstrate that recruitment of Scc2 relies on phosphorylation of H2A by Tel1 and the subsequent resection of DNA. Based on this we propose that DNA end resection affects the loading of Cohesin at DSBs therefore in two ways. First, the actual resection process to mediate the recruitment of Scc2/4. Second, the subsequent recruitment of Mec1, which enables Scc2/4 to load Cohesin at DSB. Together, these data provide a more detailed insight into the events which facilitate the recruitment of Scc2 and subsequent accumulation of Cohesin at DNA DSBs.

## Supporting information

Scherzer et al Supplementary Table 1

Scherzer et al Supplementary Table 2

## Acknowledgements

We thank Professors C. Björkegren, F. Uhlmann, J. Downs and L. Symington for strains and plasmids. This work was supported by the Swedish Research Council (2016-02206), the Swedish Cancer Society (16 0702), the Bergvall Foundation (2016-01868, 2017-02287), and the KID program at the Karolinska Institutet to L. Ström for M. Scherzer and F. Giordano.

---

**Figure S1.**
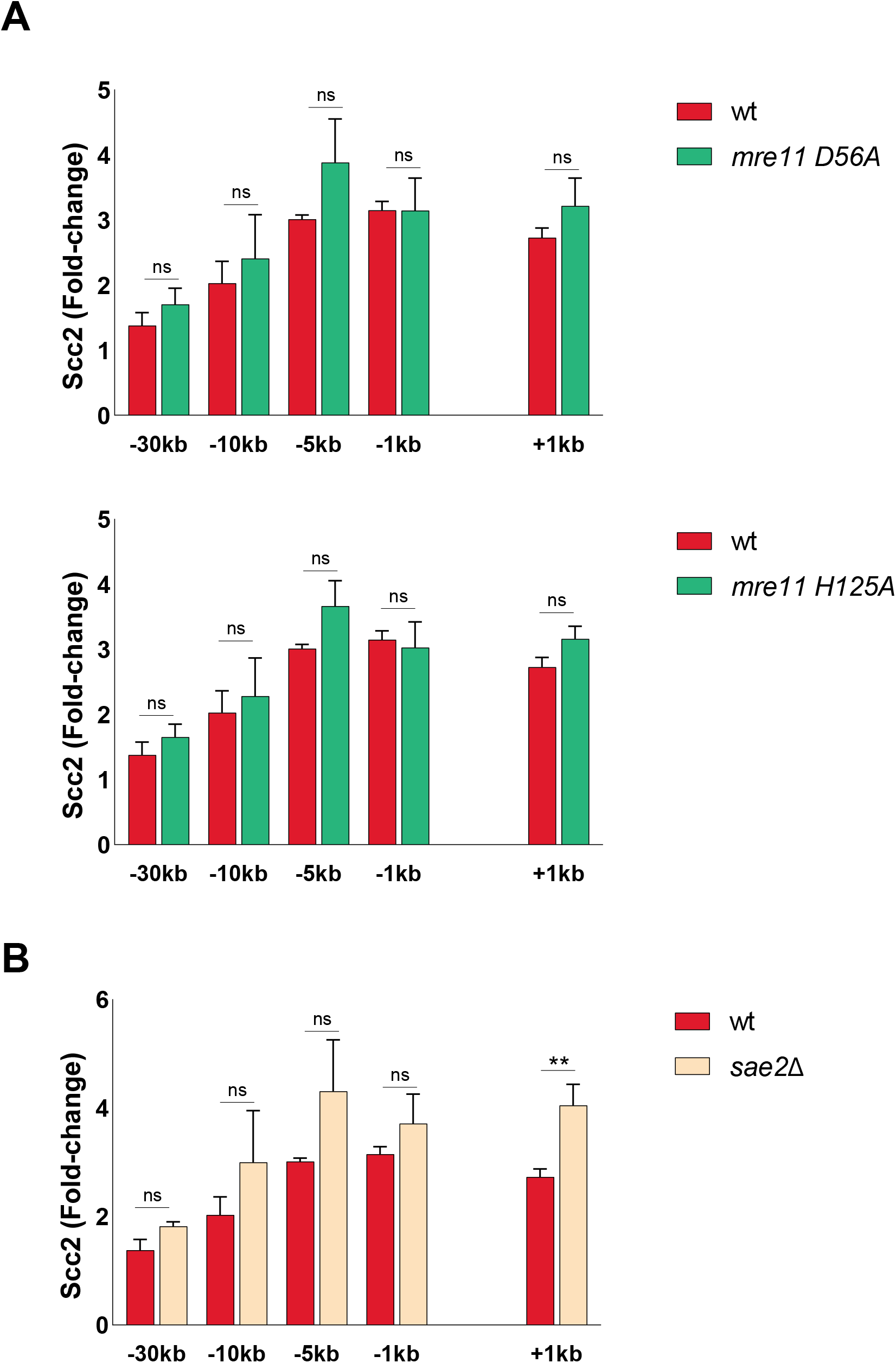
Mre11 nuclease deficiency and deletion of Sae2 do not affect Scc2 recruitment to a DSB. ChIP-qPCR of Scc2 binding at the DSB in **(a,b)** wt and **a)** a nuclease deficient *mre11 D56A* (upper) and *mre11 H125A* (lower) **b)** a *sae2Δ* strain. The graphs show means and SD of n = 3. Student’s t-test was used to compare normalized values of Scc2 between wt and indicated mutants at respective locations, 90 minutes after break induction. Significance: *p < 0,05; **p < 0,01; ***p < 0,001; ns = not significant.

**Figure S2.**
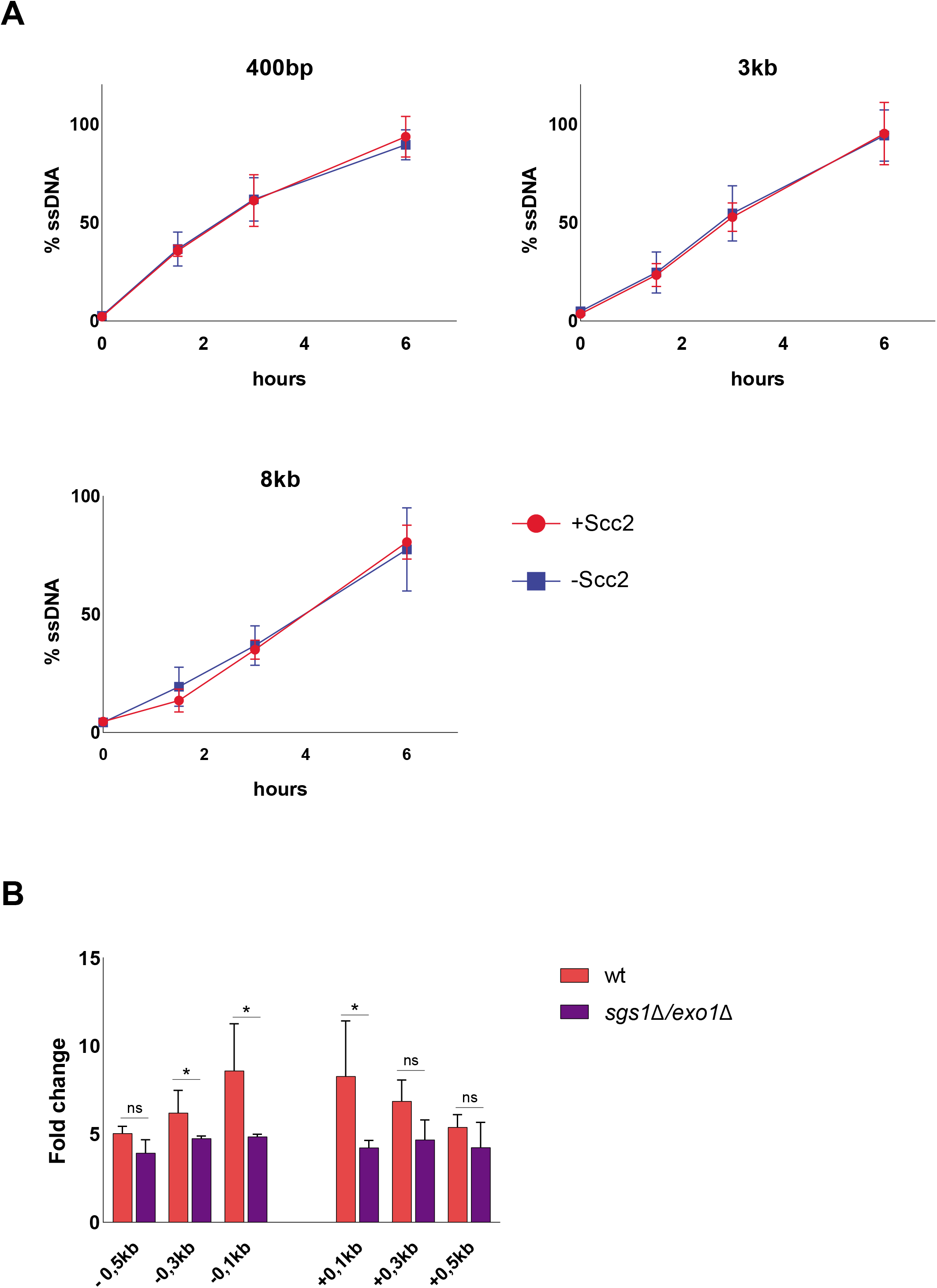
Additional information on the relationship of Scc2 and DNA end resection. **a)** Measurement of ssDNA at the DSB in the presence (red) and absence (blue) of Scc2 at indicated distances from the break. Samples were collected after addition of galactose for 0, 90, 180 and 360 minutes. The graphs show means and SD of n = 3. No significant difference was observed at any distance at any given time point using student’s t-test. **b)** ChIP-qPCR of proximal Scc2 binding at a DSB in wt and a *sgs1 Δ/exo1Δ* background. The graph shows means and SD of n = 2. Student’s t-test was used to compare normalized values of Scc2 between wt and *sgs1Δ /exo1Δ* at respective locations, 180 minutes after break induction. Significance: *p < 0,05; **p < 0,01; ***p < 0,001; ns = not significant.

**Figure S3.**
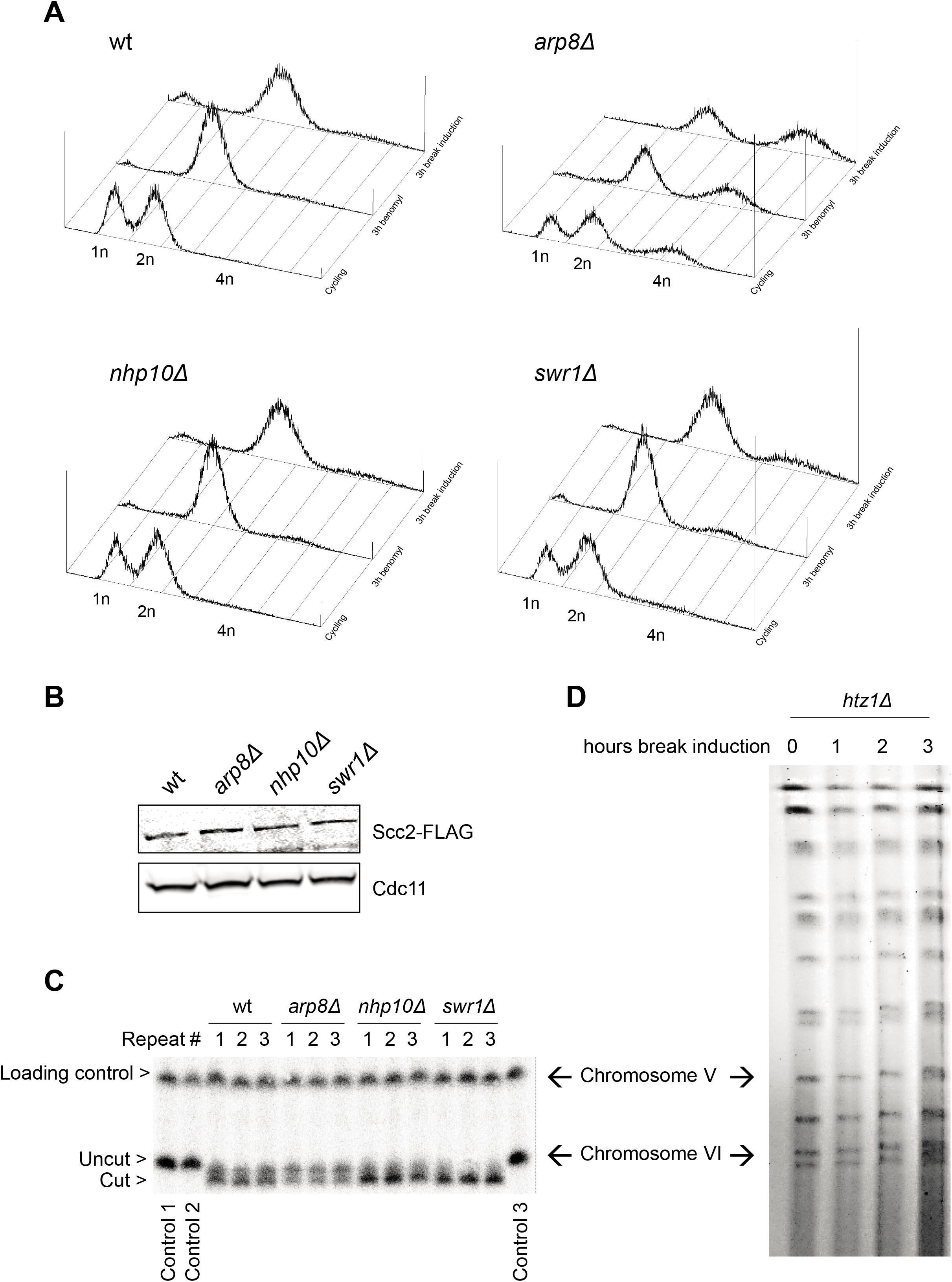
Additional information on the chromatin remodelers in our experimental setup. **a)** Representative FACS analysis showing cell cycle distribution of indicated strains at different experimental time points. **b)** Protein levels of Scc2 in cycling cells in indicated strains. **c)** Southern blot showing successful break induction of chromosome VI of indicated strains from 3 independet repeats. The probe for chromosome VI was generated using primers −1kb and −0,3kb. A probe for chromosome V was used as a loading control (see Table S2). **d)** PFGE gel showing absence of a cut chromosome VI within 3 hours of break induction (n=2).

## Notes

### Competing Interest Statement

The authors have declared no competing interest.

